# Methylation Quantitative Trait Loci are Largely Consistent across Disease States in Crohn’s disease

**DOI:** 10.1101/2020.11.16.385534

**Authors:** Suresh Venkateswaran, Hari K Somineni, Varun Kilaru, Seyma Katrinli, Jarod Prince, David T Okou, Jeffrey S Hyams, Lee A Denson, Richard Kellermayer, Greg Gibson, David J Cutler, Alicia K Smith, Subra Kugathasan, Karen N Conneely

## Abstract

**Background:** In a recent study, we identified 1189 CpG sites whose DNA methylation (DNAm) level in blood distinguished Crohn’s disease (CD) cases from controls. We also demonstrated that the vast majority of these differences were a consequence of disease, rather than a cause of CD. Since methylation can be influenced by both genetic and environmental factors, here we focus on CpGs under demonstrable genetic control (methylation quantitative trait loci, or mQTLs). By comparing mQTL patterns across disease states and tissue (blood vs. ileum), we may distinguish patterns unique to CD. Such DNAm patterns may be relevant for the developmental origins of CD.

**Methods:** We investigated three datasets: (i) 402 blood samples from 164 newly diagnosed pediatric CD patients taken at two time points, and 74 non-IBD controls (ii) 780 blood samples from a non-CD adult population and (iii) 40 ileal biopsies (17 CD cases and 23 non-IBD controls) from group (i). Genome-wide DNAm profiling and genotyping were performed using the Illumina MethylationEPIC and Illumina Multi-Ethnic arrays. SNP-CpG associations were tested via linear models adjusted for age, gender, disease status, disease subtype, estimated cell type and three genotype-based principal components. We used a Bonferroni-adjusted significance threshold to identify significantly associated SNP-CpG pairs, but also considered larger sets identified by a false discovery rate criterion

**Results:** In total, we observed 535,448 SNP-CpG associations between 287,881 SNPs and 12,843 CpG sites (*P*<8.21×10^−14^). These associations and their effects are highly consistent across different ages, races, disease states, and tissue types, suggesting that the vast majority of these mQTLs participate in common gene regulation. However, genes near CpGs associated with IBD SNPs were enriched for 18 KEGG pathways relevant to IBD-linked immune function and inflammatory responses. We observed suggestive evidence for a small number of tissue-specific associations and disease-specific ileal associations in ileum, though larger studies will be needed to confirm these results.

**Conclusion:** The vast majority of blood derived mQTLs are commonly shared across individuals. However, we have identified a subset of such, which may be involved in processes related to CD. Independent cohort studies will be required to validate these findings.

## Background

A majority of inflammatory bowel disease (IBD) trait-associated genomic variants identified in genome-wide association studies (GWAS) reside in the genome’s non-coding regions [1]. In the past decade, studies have focused on demonstrating the potential of these variants to influence specific traits or diseases through transcriptomic regulation [2-4]. Previous expression quantitative trait loci (eQTL) studies in IBD proposed that the genetic influences on gene expression are tissue-specific [5], disease-specific [6], and inflammation-specific [5, 7]. Meanwhile, recent advances in high-throughput array technologies have enabled genome-wide DNA methylation (DNAm) profiling for >850,000 CpGs. DNAm is one of the primary epigenetic mechanisms involved in gene expression regulation. Investigation of DNAm has the potential to advance our understanding of complex disease pathogenesis [8, 9]. In an epigenome-wide association study (EWAS), we recently identified 1189 CpGs showing differential DNAm in Crohn’s disease (CD) patients at diagnosis, compared to non-IBD controls [10]. A Mendelian Randomization (MR) approach revealed that most of the DNAm changes observed in CD patients are consequences of the disease rather than causal factors influencing disease risk [10]. Since DNAm can be influenced by both environmental and genetic factors, in this paper we will focus on identification of methylation quantitative loci (mQTLs), or SNPs that associate with DNAm at specific CpG sites. We will compare mQTLs found within blood across disease state and progression, as well as age and race. Finally, we will compare patterns found in blood to those found in ileal (disease) tissue. In general, we are looking for mQTL patterns that distinguish CD cases from CD controls, and mQTLs that are context-specific.

Identifying SNP-CpG associations and comparing them between disease groups will provide epigenetic annotation of IBD variants, facilitating their functional characterization. The comparative mQTL study has the potential to reveal differences in interactions between genetic and epigenetic mechanisms in IBD patients vs. controls. Further, the identification of mQTLs can aid MR approaches through identification of potential instruments to distinguish the casual vs. consequential role of DNAm in complex diseases [10], and can shed light on causal relationships between gene expression and DNAm in complex diseases [11, 12].

Investigating whether SNP-CpG associations are stable or dynamic over the course of the diseases can help inform whether the association might be a useful biomarker for disease, a useful marker for treatment, a potential cause of disease, or without obvious utility. A recent longitudinal analysis at five different stages showed that blood mQTLs are generally stable across the life span [13]. However, among the 1189 differentially methylated CpG sites identified in our previous study from blood derived DNA, we observed that during the CD disease course (at 1-3 years follow-up), DNAm patterns in cases reverted back to levels observed in healthy controls. [10]. We also noted high correlations between DNAm at these 1189 sites and levels of the inflammatory marker C-reactive protein (CRP) [10]. Investigation of mQTLs in CD patients pre- and post-treatment will reveal whether the genetic influence on DNAm at these sites varies over the disease course.

CD commonly affects the terminal ileum. Thus, in contrast to DNAm patterns in blood, methylation patterns in the ileum might be considered patterns seen at the “site of disease”. Consequently, ileal DNAm is more likely to be causative, compared to patterns observed in other tissues. Thus, we examine ileum specific mQTL associations in CD patients.

Here, we used blood samples obtained from a total of 238 pediatric subjects; 164 patients with CD and 74 healthy controls sampled from the Risk Stratification and Identification of Immunogenetics and Microbial Markers of Rapid Disease Progression in Children with Crohn’s Disease (RISK) study [14]. We replicated our analysis in an independent cohort of 780 blood samples obtained from prospectively-recruited adults in the Grady Trauma Project (GTP). Enrichment analysis showed that the previously-identified IBD GWAS SNPs are significantly enriched for mQTLs in the CD cohort, but not in the adult cohort. To assess whether the mQTLs are specific to disease, disease course, or tissue type, we performed separate mQTL analyses in 164 CD cases at diagnosis, in cases at follow-up (1-3 years from the day of diagnosis), in 74 healthy controls and in 40 ileal biopsies (obtained from a subset of subjects included in the blood mQTL analysis; 23 controls and 17 patients with CD at diagnosis).

## Results

### Blood mQTL discovery

To identify blood mQTLs, we tested the association between 3,109,863 SNPs and 609,192 CpG sites from 238 blood samples taken from 164 pediatric patients with CD at diagnosis and 74 matched non-IBD controls (Additional file 1: Table S1). We identified 535,448 associated SNP-CpG pairs involving 303,986 (9.8%) unique SNPs and 13,823 (2.3%) unique CpGs, at a Bonferroni adjusted genome-wide significance of *P* < 8.21 × 10^−14^ (Fig. 1A). On average, each CpG site was associated with ∼38 SNPs (interquartile range (IQR): 4 – 42), which likely reflects linkage disequilibrium (LD) between proximal SNPs, while each mQTL variant associated with 1.73 CpGs (IQR: 1 to 2). Using the *clump* command in PLINK [15], we identified 16,313 LD pruned proximal SNPs that were associated with 13,202 CpG sites, which made a total of 24,523 associations.

**Fig. 1:**
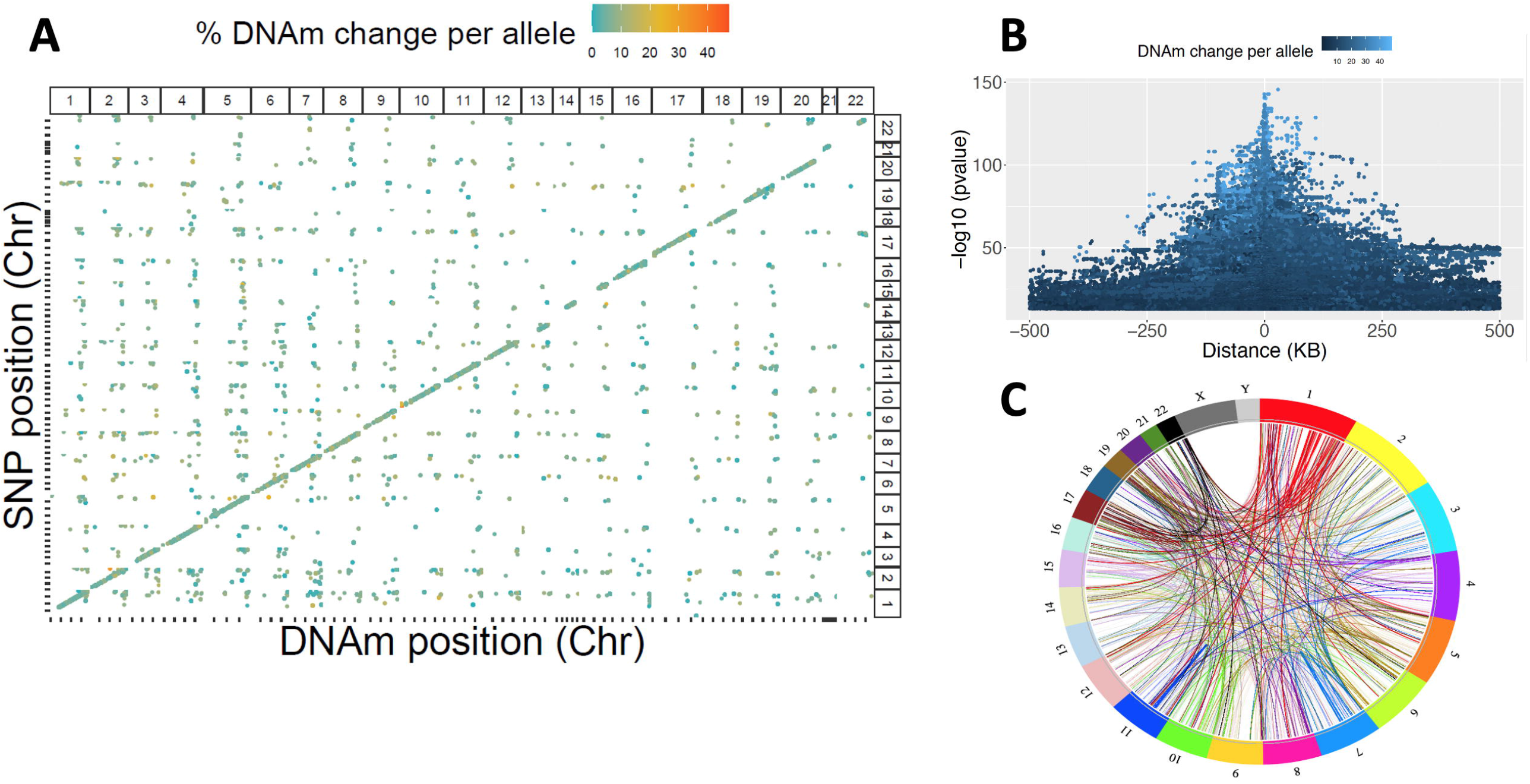
Genetic associations with DNAm genome-wide in discovery cohort. (A) In total 535,448 SNP-CpG pairs are plotted, including both *cis*- and *trans*-associations. The X-axis indicates genomic variant position, whereas the Y-axis indicates CpG site position. The color of each point represents absolute differences in DNAm per allele relative to the reference allele. Point size represents the significance of the association. A clear positive diagonal line indicates that the majority of associations are *cis* associations. (B) 498,261 *cis* associations are plotted by position of CpG relative to SNP. The strongest associations are seen for proximal pairs of SNPs and CpG sites. (C) 37,187 *trans-*associations are plotted, with most of them occurring within the same chromosome.

93% (n=498,261) of the SNP-CpG pairs involved associations between a genetic variant and CpG site within 1 MB (±500 kb; Fig. 1B) and were considered *cis-*acting. The remaining 7% appeared to be *trans-*acting (n=37,187); of these, 44% (n=16,457) involved CpGs and SNPs from the same chromosome that were >500 kb apart, while 56% (n=20,730) involved a SNP and CpG from different chromosomes (Fig. 1C). On average, *cis*-acting mQTLs had larger effects with a mean effect size of 8.7% (IQR: 5.3 – 10.8) change in DNAm per allele, compared to *trans*-acting mQTLs with a mean effect size of 6.9% (IQR: 2.8 – 9.4) change in DNAm per allele (Mann-Whitney-Wilcoxon *P* < 2.2 × 10^−16^. (Figure S1). Similarly, on average cis-acting mQTLs had larger effect sizes, with a mean effect size of 0.0046 (IQR: -0.06 – 0.07; SD: 0.097), compared to trans-acting mQTLs, with a mean effect size of 0.0035 (IQR: -0.06 – 0.053; SD: 0.081) (Figure S1). A list of all *cis-* and *trans-*SNP-CpG pairs identified from 238 blood samples is provided in Additional file 2: Table S2. Enrichment analysis of cis-mQTL associated SNPs (n=287,881) showed that they are more likely than other SNPs on the array to occur 1 kb upstream of transcription start site (TSS), 1 kb downstream of transcription end site (TES), in exons, the 3’ or 5’ untranslated regions, or within 2 bp of a splicing junction (*P*<2×10^−10^; Additional file 1: Table S3). Similarly, cis-mQTL associated CpGs (n=12,843) are more likely than other CpGs on the array to occur in CpG islands and shores (*P* < 2.2 × 10^−16^; Additional file 1: Tables S4-5.)

### Blood mQTLs are highly replicable in an independent dataset regardless of age and race

To evaluate whether our blood mQTLs are unique to IBD, or more common to the human condition, we tested all *cis* associations in an independent cohort of 780 blood samples (GTP cohort) collected previously [16]. More details about demographic information of this dataset are available in Almli *et al*. [16] as well as in our methods section. Notably, while our discovery dataset consists of pediatric patients with predominantly Caucasian ancestry (78%), the GTP dataset is composed of prospectively-recruited adult individuals with predominantly African American ancestry (93%), with a mean age of 42.26 (SD: 12.31) years (Additional file 1: Table S1). After QC, 5,971,966 genomic variants and 608,245 CpG sites were retained for 780 blood samples. To test for the replicability of blood mQTLs from the discovery cohort, we focused only on the SNPs and CpG sites identified in the discovery cohort. In total, 253,000 out of 287,881 SNPs (87.9%) and 12,808 out of 12,843 CpG sites (99.7%) were available to be tested in the replication cohort, representing a total of 425,288 CpG-SNP relationships. Applying a standardized genome-wide significance threshold of *P* <5 × 10^−8^, we replicated ∼84% of the CD dataset associations (358,804 SNP-CpG pairs out of 425,288 tested). This includes 87% (n=220,468) of SNPs, and 85% (n=10,905) of CpG sites. At a less stringent nominal significance threshold of *P* <0.05, we detected 388,609 associations, of which 91.3% (n=388,217) were considered replications, where the effect sizes of both CD and GTP cohort associations are directionally consistent. Test statistics for these associations were highly correlated in the discovery vs. replication cohorts (R^2^=0.94, *P* < 2.2 × 10^−16^; Fig. 2), indicating that most blood SNP-CpG associations are consistent regardless of age, race and disease status. Summary statistics for the 388,609 SNP-CpG associations are provided in Additional file 3: Table S6. Among the few mQTLs showing associations in the opposite direction (n=393; 358 unique SNPs and 119 CpG sites) between the CD and GTP cohorts (i.e. a SNP associated with increased methylation in one cohort, and decreased methylation of the same CpG in another cohort), we did not observe significant enrichment for any KEGG pathways, including inflammation or immune-related pathways (data not shown). Similarly, none of the previously identified IBD-associated SNPs [17, 18] or CD-associated CpGs [10] were involved in these 393 mQTL associations. These results are consistent with our previous [19] findings that blood mQTLs are stable across ancestries (African American vs. Caucasian), and developmental stages (neonate vs. adult). Collectively, our results indicate that the vast majority of genetic influence on changes in blood DNAm are highly consistent regardless of disease, race and age groups, and thus unlikely to be casually related to CD in any substantial way.

**Fig. 2:**
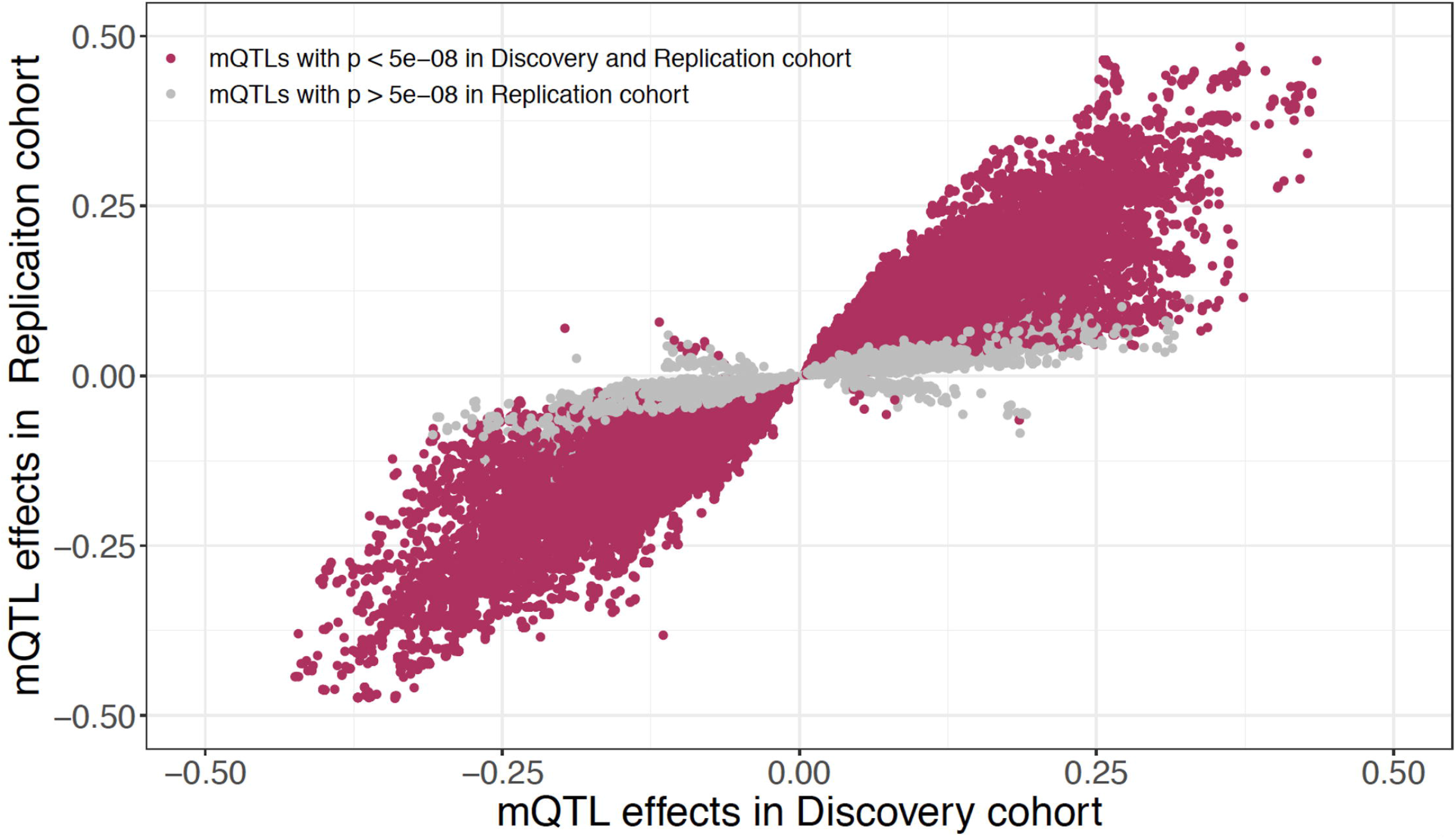
Replication of mQTL associations in replication cohort. Effect sizes for 425,288 SNP-CpG pairs are compared between discovery and replication cohorts. Associations significant in both discovery cohort (*P* < 8.21 × 10^−14^) and replication cohort (*P* < 5 × 10^−8^) are marked in maroon. Gray dots indicate associations significant only in discovery cohort.

### CD-specific mQTLs in blood samples

To assess whether mQTLs vary with respect to disease status, we retested each of the 498,261 significant SNP-CpG pairs separately within controls (n=74) and within CD cases (n=164). We observed a strong positive correlation (R^2^=0.97, *P* < 2.2 × 10^−16^) between effect sizes estimated from controls vs. cases (Fig. 3A). Collectively, our data establish that while methylation levels may vary with CD at specific sites (CD-associated CpG sites) [10], the influence of SNPs on CpG methylation levels at CpG sites remains consistent in the presence or absence of disease. Results obtained for CD cases and healthy controls are provided in Additional file 4: Table S7.

**Fig. 3:**
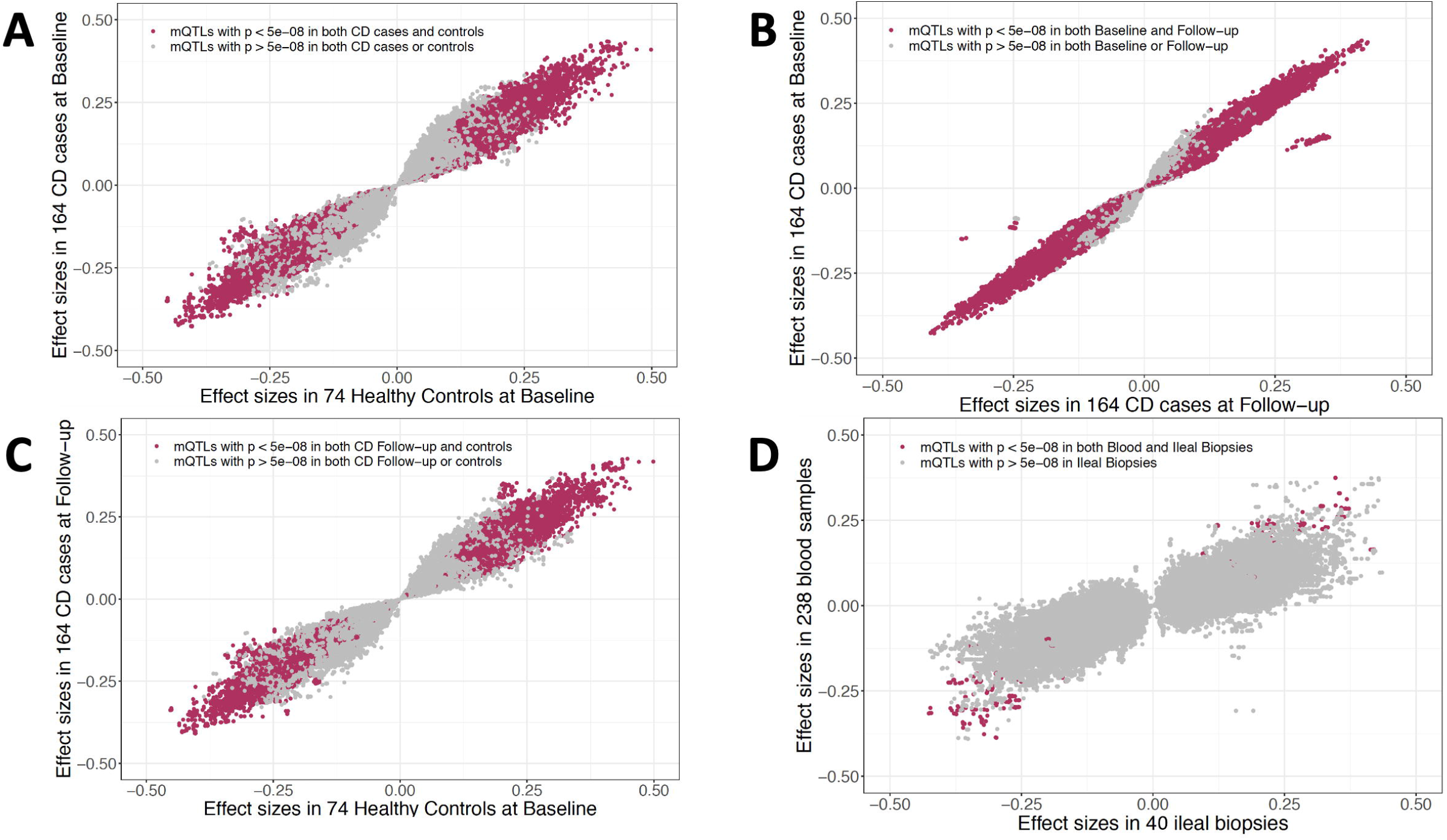
Disease-, disease course- or tissue-specific associations in discovery cohort. For 498,261 *cis* SNP-CpG pairs significant in the discovery cohort of 238 blood samples, Figure 3 shows effect sizes estimated in the following subgroups: (A) 74 healthy controls vs. 164 CD cases at diagnosis, (B) 164 CD cases at diagnosis vs. their 3-year follow-up, (C) 74 healthy controls at baseline vs. 164 CD cases at follow-up, and (D) all 238 blood samples vs. 40 ileal biopsies. Maroon color indicates the associations that are significant (*P* < 5 × 10^−8^) in both subgroups. Gray indicates the associations that are significant in one subgroup but not the other.

### Longitudinal profiling of mQTLs

In our previous report, we demonstrated that with treatment, DNAm changes associated with CD at diagnosis revert to the levels seen in healthy controls [10]. To examine the longitudinal trajectory of SNP-CpG associations and their dynamics during the course of the disease, we took advantage of our longitudinal DNAm data generated from same individuals from the discovery cohort (164 CD cases at diagnosis vs. follow-up). We found that mQTL effect sizes were similar between diagnosis and at 3-year follow-up in CD patients, showing a strong positive correlation in magnitude and direction of effect (R^2^=0.99, *P* < 2.2 × 10^−16^; Fig. 3B; Additional file 4: Table S7). All discovery SNP-CpG associations (n=497,689) were nominally significant at baseline (*P* < 0.05), and 99.5% (n=497,443) were nominally significant during the course of the disease (*P* < 0.05). Therefore, our results suggest that these genetic associations are independent of disease status or treatment.

### Tissue-specific associations between blood and ileum

To further evaluate how the mQTLs identified in our blood-based discovery dataset act in tissues more directly relevant to disease, we estimated SNP-CpG effect sizes using DNAm data from ileal biopsies (n=40) and compared them to effects observed in blood. With the standard replication threshold of *P* < 0.05, we replicated 50.4% (n=238,400) blood SNP-CpG associations in ileal biopsies. Overall, we noted an extremely strong correlation in effect sizes between ileal and blood samples, with 91.4% (n=473,382) of mQTLs showing directional consistency (R^2^=0.83, *P* < 2.2 × 10^−16^; Fig. 3D). A list of all ileal biopsy mQTL results are provided in Additional file 5: Table S8. We further investigated CpGs (n=2,188) from SNP-CpG associations (n=24,879) with directionally inconsistent effects between blood and biopsy for tissue-specific biological functions, and observed that these sites were more likely to localize to genes involved in metabolic pathways (*P* = 3.34 × 10^−8^; Table S9). Using summary statistics from two different studies, Qi et al [20] recently showed that mQTLs are highly correlated among the blood and brain tissues. Similarly, using blood and four regions of postmortem brain samples, we previously showed a strong mQTL correlation among the tissues, especially an extreme overlap within the four brain regions [19]. Our current results show a similar pattern for blood vs. ileum taken from the same patients, where the mQTL effects are highly correlated between blood and ileal biopsies.

### Enrichment of previously-identified IBD susceptible SNPs in blood mQTLs

To date, large-scale GWAS meta-analysis have identified 241 susceptibility loci for IBD [17, 18]. A majority of these (>90%) localize to non-coding regions, presenting a key challenge to identify the functional variants and the relevant genes they act upon. Of these 241 previously-identified IBD SNPs, 168 were genotyped in our discovery dataset. We found that 37 (22.2%) of these 168 SNPs associated with 69 CpG sites in our discovery set of mQTL associations (*P* < 8.21 × 10^−14;^ Figure S2A; Additional file 6: Table S10). Notably, all of these associations were cis-mQTL associations: 90% (n=62) of these CpGs were located within 100 kb from the associated GWAS variant. Of these 62 CpGs, 13% are located within 1.5 kb of a transcriptional start site (TSS200 or TSS1500), and 47.8% are between the start and stop codon. These results provide an additional 31 genes potentially associated with the previous-identified IBD SNPs, where the associated CpGs may be located within the gene body or near the TSS or promotor regions of the gene. Notably, 4 of the established IBD-associated SNPs (rs13407913, rs798502, rs59043219 and rs1819333) showed association with >5 CpGs located within the gene bodies of *ADCY3, GNA12, IRF6*, and *RNASET2/RPS6KA2*, respectively. The 37 IBD-associated SNPs comprised .013% of the total number of mQTL SNPs observed in our discovery cohort (Additional file 6: Table S10). For comparison, in our GTP cohort, which did not include IBD cases, we observed associations between 87 IBD-associated SNPs and 252 nearby CpGs. However a greater number of mQTL associations were observed in total in this larger cohort, the 87 IBD-associated SNPs comprised a smaller percentage of the total (0.006%; Additional file 6: Table S10). This suggests that the mQTLs identified in our discovery cohort (which is 69% IBD patients) are significantly enriched IBD-associated SNPs (OR = 2.1 [CI: 1.4 to 3.1], Fisher’s exact *P* = 2.71 × 10^−4^). Summary statistics for 69 and 253 SNP-CpG associations from the IBD susceptibility SNPs identified in both discovery and replication cohorts are provided in Table 1.

We further searched for previously-identified IBD SNPs in a wider set of mQTL results identified using a less stringent significance threshold (FDR<0.05) in the discovery cohort. At this threshold, 64.2% (n=108) of IBD SNPs were associated with 453 CpGs within 1 MB, comprising 454 cis-mQTL associations (Additional file 6: Table S10). Of these CpGs, 84.5% (n=384) were located within 100 kb of the associated GWAS variant (Fig. 4A). Further disease-specific analysis on these mQTLs showed that similar to SNPs across the genome, associations between IBD-associated SNP and DNAm in blood are extremely consistent between CD cases (n=164) vs. controls (n=74) (Fig. 4B, Additional file 6: Table S10).

**Fig. 4:**
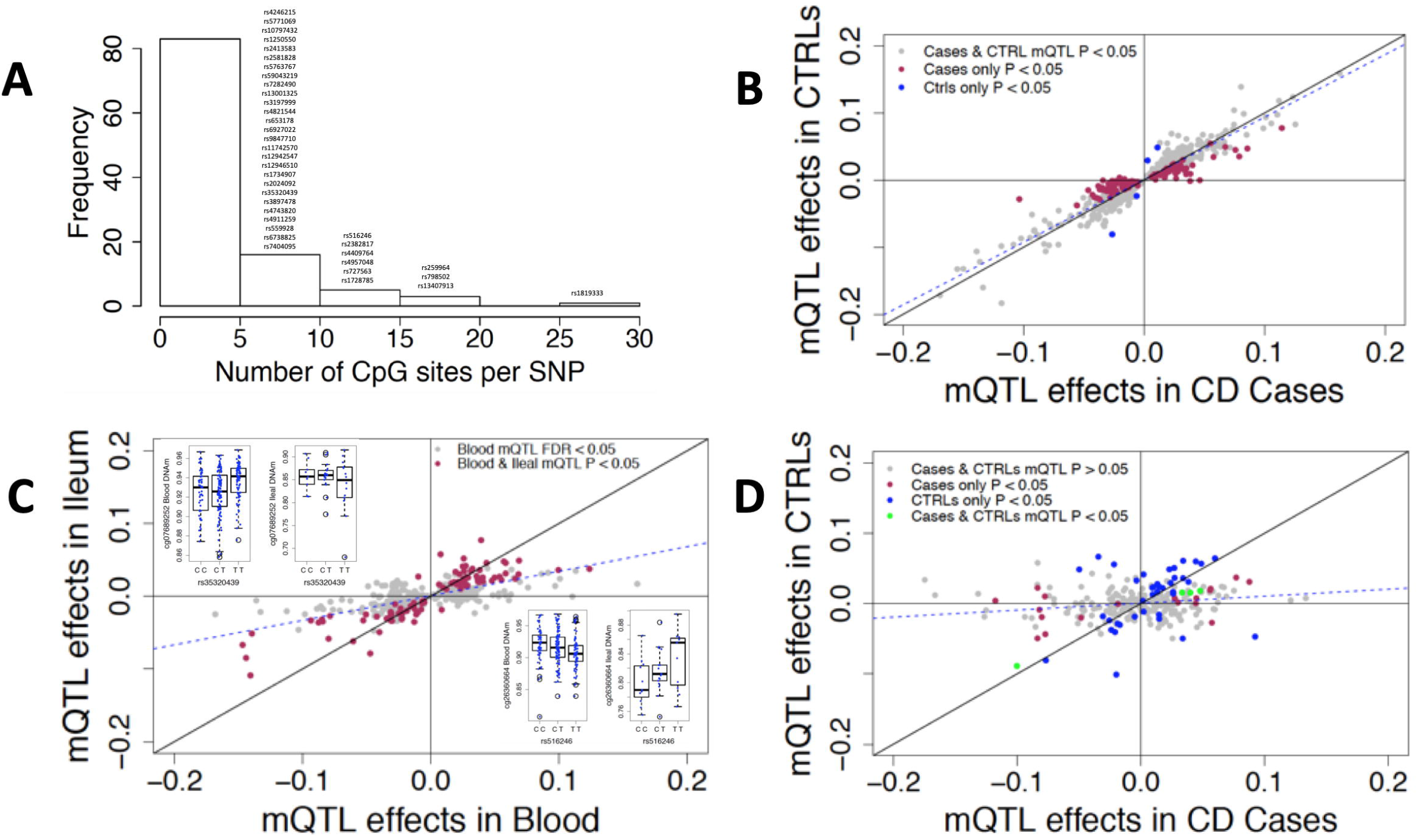
Comparing CpG-specific associations with IBD SNPs in blood vs. ileum. (A) Frequency of CpG sites that are associated with previously-identified IBD SNPs in discovery cohort at FDR < 0.05. In total, 454 associations are considered. (B) Effect sizes of 454 SNP-CpG associations compared in CD cases vs. controls in blood. X-axis shows the effect sizes in CD cases (n=164) and Y-axis shows the effect sizes for the same associations in controls (n=74) (C) Effect sizes of 370 SNP-CpG associations found in blood are compared to effect sizes observed in ileum. The nominally significant associations in ileum (*P*< 0.05) are marked in maroon. (D) Effect sizes of 269 SNP-CpG associations compared across the disease groups in ileum. X-axis shows effect sizes in CD cases (n=17) and Y-axis shows effect sizes for the same associations in controls (n=23).

### Pathway enrichment of CpG sites that are under the influence of IBD-associated SNPs

To investigate the utility of DNAm as a tool to identify functional variants and underlying regulatory mechanisms that may contribute to our fundamental understanding of CD susceptibility and pathogenesis, we examined whether CpG sites associated with IBD-associated SNPs localize near genes associated with specific pathways. We first evaluated all 69 CpGs whose blood DNAm associated with IBD SNPs in the discovery analysis (*P* < 8.21 × 10^−14^) to identify the novel pathways that are relevant to CD. Our pathway enrichment analysis identified 18 KEGG pathways that were more likely to occur in the IBD SNP– associated CpGs than would be expected by chance (FDR < 0.05; Additional file 1: Table S11). Among these, five were relevant to immune function and other pathways, including human cytomegalovirus infection, insulin resistance, vascular smooth muscle contraction, hepatitis B, and chemokine signaling. Pathway enrichment analysis on an expanded set of CpG sites (n=454) that associated with IBD SNPs according to a looser significance threshold (FDR < 0.05) showed enrichment for 7 KEGG pathways (FDR < 0.05), including a natural killer cell mediated toxicity pathway. We also observed nominally significant (*P* < 0.05) enrichment for immune and inflammatory related pathways such as MAPK signaling pathway, cytokine-cytokine receptor interaction, inflammatory bowel disease (IBD), JAK-STAT signaling pathway, and Th17 cell differentiation (Additional file 1: Table S12).

### Influence of previously-identified IBD SNPs on DNAm in ileum mQTLs

We next investigated the 37 SNPs that were both IBD-associated [17] and associated with CpGs in our discovery cohort (*P*<8.21×10^−14^) to identify whether they were also mQTLs in ileum. Seventeen of 37 SNPs were nominally significant mQTLs (*P* < 0.05) in ileum as well, and effect sizes showed a strong positive correlation between ileum and blood (R^2^ = 0.789, *P* <7.87 × 10^−16^, Figure S2C; Additional file 7: Table S13).

We also followed up on the 108 FDR-significant IBD-associated SNPs in ileum. We tested 370 of the 454 blood mQTL pairs for association in ileum. These mQTL pairs included 85 of the 108 IBD-associated SNPs; the remaining 23 SNPs were not tested due to insufficient genetic variation in the smaller number of ileum samples. At a nominal threshold of *P* < 0.05, we detected associations for 51 of 85 IBD-associated SNPs. In total, 91 (24.6%) of 370 blood mQTLs in ileum were nominally significant, and the effect sizes were consistent in direction for 89 of these, indicating replication (Fig. 4C; Additional file 8: Table S14). The two IBD SNPs showing association in the opposite direction in ileum (rs35320439, and rs516246) were independently associated with CpGs cg07689252 and cg26360664 (Fig. 4C). Neither of these CpGs were CD-associated CpGs. The strong positive correlation and directionally consistent effects (R^2^=0.87; *P* < 2.2 × 10^−16^; Fig. 4C) suggest that the IBD-associated SNPs influence the DNAm levels mostly similarly across both tissues. Notably, performing this analysis separately in CD cases (n=17) and controls (n=23) in ileum showed disease-specific patterns for 13 of the IBD associated SNPs (Fig. 4D; Additional file 8: Table S14). These results suggest there may be disease-specific SNP-CpG associations in actual disease-relevant tissues such as ileum for CD. However, given our smaller sample size (n=40) and the nominal significance level, further confirmation is needed in larger studies of disease-relevant tissues such as ileum.

### CD-associated differentially methylated CpGs in blood mQTLs

Recently, we identified 1189 CD associated CpG sites whose methylation level in blood distinguished CD cases from non-IBD controls [10]. After exclusion of 212 CpGs with one or more SNPs in their probe sequence, we tested 977 of our previously-identified CpG sites for associations with SNPs within 1MB. In the discovery cohort, 22 (1.8%) of these 977 CpGs associated with 565 unique SNPs for a total of 565 SNP-CpG associations (*P* < 2.81 × 10^−14^; Figure S2B; Additional file 9: Table S15). Although we did not observe an enrichment of CD-associated CpG sites compared to the replication cohort (Table 1), one CD-associated CpG site cg20406979 had a strong association (*P* = 7.60 × 10^−25^) with SNP rs1819333, previously reported to associate with CD [10]. This SNP and CpG site were located 314 nucleotides apart, 3kb upstream of the transcription start site for the gene *RNASET2*.

We further investigated the 977 CD-associated differentially methylated CpGs using an FDR-based significance threshold (FDR<0.05) and observed a total of 7819 cis-associations. 29% (n=285) of 977 CD-associated CpGs were associated with 7,120 nearby genomic variants (Additional file 9: Table S15), with 59.9% of these CpGs associating with more than 5 SNPs (Fig. 5A). Further separate analysis on CD cases (n=164) and controls (n=74) indicates that the mQTLs associated with CD-associated DNAm sites in the blood are extremely consistent across the disease types (Fig. 5B; Additional file 9: Table S15).

**Fig. 5:**
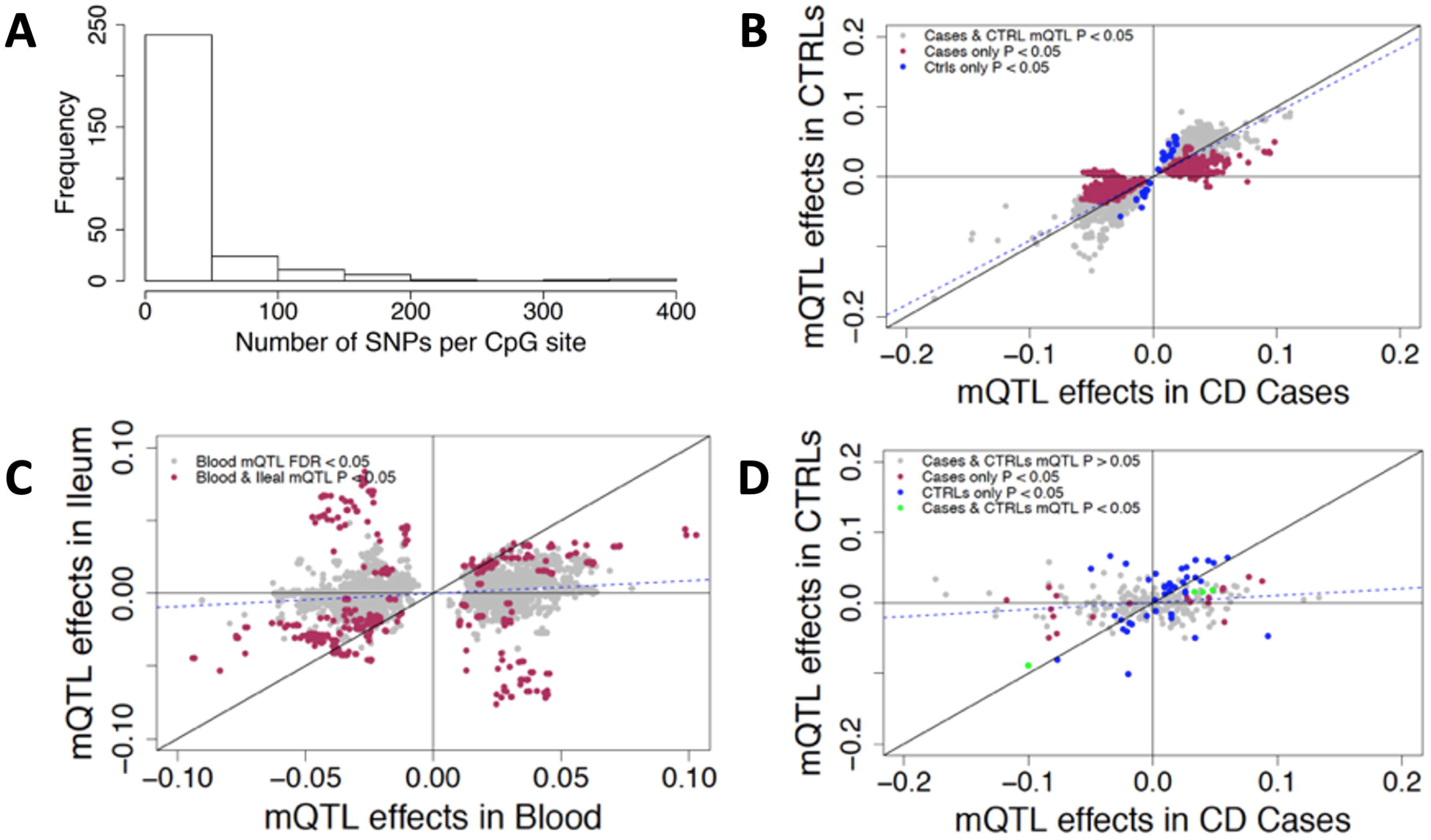
Testing the effects of CD-associated CpG sites in blood and ileum. (A) Frequency of SNPs that are associated with CD-associated CpG sites in discovery cohort at FDR < 0.05 are plotted. In total, 7819 associations with 289 CD-associated CpG sites are considered. (B) Effect sizes of 7819 SNP-CpG associations compared in cases vs. controls in blood. X-axis shows the effect sizes in CD cases (n=164) and Y-axis shows the effect sizes for the same associations in controls (n=74) (C) Effect sizes of 6221 SNP-CpG associations from blood are compared to effect sizes observed in ileum. Associations nominally significant in ileum (*P* < 0.05) are marked in maroon. (D) Effect sizes of 4142 SNP-CpG associations compared in cases vs. controls in ileum. X-axis shows effect sizes in CD cases (n=17) and Y-axis shows effect sizes for the same associations in controls (n=23).

Next, we investigated whether the CD-associated CpGs associated with 108 previously-identified IBD risk SNPs within 1MB. We observed 2 IBD SNPs (rs1819333, rs3853824) associated with 4 CD-associated CpGs (cg20406979, cg23216724, cg15706657, and cg15815084) for a total of 4 cis associations. Of those, three CpGs cg20406979, cg23216724, and cg15706657 had stronger association with one IBD associated SNP rs1819333 (*P* = 7.60 × 10^−25^, *P* = 1.2 × 10^−11^, & *P* = 9.3 × 10^−07^, respectively; Additional file 9: Table S15).

### Influence of SNPs on CD-associated CpGs in Ileum mQTLs

Next, we investigated 22 of the 1189 CpGs previously associated with CD in blood [10] that were also blood mQTLs (*P* < 8.21 × 10^−14^) to see if they are also mQTLs in ileum. In blood, these CpGs were involved in 565 significant SNP-CpG associations. We replicated 173 (30.6%) of these associations in ileal biopsies using a nominal threshold of *P* < 0.05, and observed effect sizes are consistent in magnitude and direction in ileum vs. blood (R^2^ = 0.82; *P* < 2.2 × 10^−16^; Figure S2D; Additional file 10: Table S16).

We performed similar comparisons for the 285 CD-associated CpGs that were significant at FDR<.05 in blood, which were involved in 7819 significant SNP-CpG associations in blood. We followed up on 6,227 of these SNP-CpG pairs in ileum; the remainder were not tested due to insufficient genetic variation among the 40 ileal samples. At a nominal threshold of *P* < 0.05, we detected 47 of 285 significant CD-associated CpGs and 1,002 (16.1%) out of 6,227 blood SNP-CpG pairs in ileum (P < 0.05). Among these 1002, 65.4% (n=655) showed directionally consistent associations (R^2^ = 0.92; *P* < 2.2 × 10^−16^; Fig. 5C). The 34.6% (n=347) of CpG-SNP associations that showed association in the opposite direction in blood vs. ileum (Additional file 10: Table S16) comprised 12 CD-associated CpGs that were associated with 341 SNPs. Boxplots for the top SNP associated to the 12 CD-associated CpGs are compared between blood and ileum (Additional file 10: Table S16). None of the IBD SNPs showed the opposite mQTL effects between blood and ileum. Overall, only one IBD SNP (rs1819333) was nominally associated with a CD-associated CpG (cg15706657) in ileum (*P* = 0.040), and the association was directionally consistent with blood results in our discovery cohort. Our analysis stratified by CD suggested possibly distinct effects for cases (n=17) vs. controls (n=23) in 120 mQTLs (115 SNPs and 13 CD-associated CpGs) in ileum (Fig. 5D, Additional file 11: Table S17), though a much larger sample would be required to test this formally.

## Discussion

The primary focus of this study was to identify and characterize mQTLs in blood from 238 pediatric samples of European ancestry. Our findings establish that blood mQTLs are robust and reproducible, supporting their generalizability across age groups, ancestries, disease status, and DNA sample source, and offer a valuable source for future studies examining blood mQTLs. Recent studies have identified different sets of differential DNAm patterns in various tissue types [21, 22], and previous epigenomic and transcriptomic studies in IBD suggest that disease-relevant tissue samples are needed to develop precise biomarkers for the disease subtypes [5, 23-25]. Our findings show that among genome-wide-significant mQTLs, genetic influences on DNAm are generally consistent across blood and ileum, even among IBD SNPs and CD-associated CpG sites. However, we observed lower cross-tissue reproducibility using a relaxed FDR-based significance criterion. Using a threshold of FDR < 0.05 in blood and nominal association (P < .05) in ileum, we identified two IBD SNPs (Figure 4C) and 12 CD-associated CpGs (Figure 5C; Figure S3) with associations in the opposite direction between blood and ileum. This could suggest that a small portion of mQTLs act in a tissue-specific manner, but could also be due to the low power and relaxed significance criteria in the ileal analysis. Studies with large numbers of samples from both blood and ileal biopsies from the same cohort are needed to formally test for tissue-specific associations for the previously-identified IBD SNPs or CD-associated CpGs.

Blood mQTLs were also consistent in comparisons of CD patients pre-vs. post-treatment and in CD patients compared to healthy matched controls or to an external epidemiologic sample of predominantly African American adults. This trend was observed even when we focused on previously-identified IBD SNPs and CD-associated CpGs, though was not as clear when studied in the ileal biopsies, again likely due to the small numbers.

Our study has several limitations. We did not explore associations involving rare variants in this study given limited power to identify such associations. In particular, because of the limited number of ileal samples used in this study, power was limited to identify tissue-specific mQTLs or to detect functionally important associations involving previously-identified IBD variants in the disease-relevant ileum.

## Conclusions

We performed the first large mQTL study in the IBD research community that allows comparisons pre- and post-treatment, between disease groups, and between blood and ileum sampled from the same patients. The vast majority of blood derived mQTLs are commonly shared across individuals. Also, our study showed a smaller portion of tissue- and disease-specific SNP-CpG associations for previously-identified IBD-associated SNPs or CD-associated CpGs in the blood. In contrast, such associations in ileum are not clear, warranting further study in disease relevant tissue samples from a larger cohort.

## Methods

### Study Population and design of discovery cohort

The discovery cohort was a subset of pediatric subjects recruited under the Risk Stratification and Identification of Immunogenetic and Microbial Markers of Rapid Disease Progression in Children with Crohn’s Disease (RISK) study [14] for whom DNAm data had been collected. A detailed description of the dataset is provided in our previous study [10]. In total, 164 newly diagnosed, pediatric patients with Crohn’s disease (cases) at two time-points (diagnosis and 3-year follow-up) and 74 non-IBD controls had DNAm measured for this study. Patients with no bowel pathology upon endoscopy, negative gut inflammation, and continued presentation as asymptomatic for IBD during follow-up were considered as non-IBD controls.

### Blood genotype data for CD cohort

Peripheral blood DNA genotypes were obtained for 238 subjects using Infinium Multi-Ethnic Global-8 Kit (Illumina, San Diego, CA) and processed with GenomeStudio software. Detailed quality control (QC) steps for both samples and variants are provided in our earlier study [10]. Briefly, sample QC was performed by checking that (i) genotype call rates >95%, (ii) inferred gender is consistent with clinical records, and (iii) subjects are unrelated according to a pairwise identity-by-descent test. Similarly, SNP QC was performed by identifying and removing SNPs with low call rate (<95%), (ii) Hardy-Weinberg disequilibrium (*P*<1.0×10^−3^), or (iii) minor allele frequency (MAF) < 5%, as well as (iv) non-autosomal variants and (v) variants mapping to multiple locations. This resulted in a discovery data set consisting of 636,006 high-quality variants. These variants for all 238 subjects were then used to impute genotypes from the 1000 Genomes Project Phase3 autosomal reference panel, using IMPUTE2 software [26]. Imputed SNPs with MAF<1% and imputation quality scores <80% were removed. We also removed imputed variants with no dbSNP annotation, INDELs/CNV, and SNPs with missing genotype information in >5% of the samples. Lastly, SNPs with <5 individuals in each group (homologous reference (AA), heterozygous (AB), and homologous variant (BB)) were excluded. In total, 3,109,863 high-quality SNPs for all 238 subjects were retained across the entire genome.

### Blood and Ileal DNA methylation data for CD cohort

DNAm data was profiled for 402 samples (164 CD cases at diagnosis and 3-years follow-up and 74 healthy controls at diagnosis) as described in our previous study [10]. In this study, we have added DNAm data for 40 ileal biopsies, performed in 17 CD cases and 23 non-IBD controls from the discovery cohort. Genome-wide DNA methylation for these samples was profiled at single-base resolution using MethylationEPIC BeadChips. QC for the blood DNA methylation is described in detail in [10]. We applied the same QC protocol on the ileal biopsies and retained all 40 ileal biopsies. Data was normalized using the beta-mixture quantile dilation (BMIQ) method [27]. Estimated cell counts for CD4+ T cells, CD8+ T cells, NK cells, B cells, monocytes, and granulocytes were calculated using Houseman’s approach [28]. After exclusion of CpGs with SNPs in their probe sequences, DNA methylation of 609,192 CpG sites in 402 samples from 238 subjects was used in this study.

### Blood genotypes for GTP cohort

Study design and genotyping of the replication cohort was previously described in recent studies [29, 30]. Briefly, GTP individuals were prospectively recruited from waiting rooms at Grady Memorial Hospital in Atlanta, GA between 2005-2008 for a study of stressful life events and symptoms of post-traumatic stress disorder (PTSD). 791 subjects from GTP were genotyped using Illumina HumanOmni1-Quad, HumanOmniExpress or Multi-Ethnic Global arrays. Genotypes were called using Illumina’s GenomeStudio software. QC analyses on the genetic data from each chip were done separately, and untyped variants were imputed with IMPUTE2 software [26] using 1000 Genomes Project Phase3 as a reference panel. Individuals with > 5% missing data were removed. SNPs with > 95% call rate, MAF > 1% and Hardy–Weinberg equilibrium (HWE) *P* > 10^−5^ were included. As a result, 5,971,966 genomic variants for 780 samples were retained. We used PLINK software [15] to perform genotype QC in both discovery and replication cohorts.

### Blood DNAm for GTP cohort

DNA was extracted from whole blood and interrogated using the MethylationEPIC BeadChip according to manufacturer’s instructions. Raw methylation beta values from the Infinium MethylationEPIC BeadChip were determined via the Illumina GenomeStudio program. Samples with probe detection call rates <90% and those with an average intensity value of either <50% of the experiment-wide sample mean or <2,000 arbitrary units (AU) were removed using the R package CpGassoc [31]. Data points with detection p-values >0.01 were set to missing. CpG sites that cross-hybridize between autosomes and sex chromosomes were removed. BMIQ was used to normalize the distribution of types I and II probes [27]. In total, methylation of 608,245 CpG sites was examined in 780 blood samples of the replication cohort.

### mQTL (Methylation Quantitative Trait Loci) study

All SNPs passing QC were tested for association with ∼600K CpG sites using a linear mixed model implemented in matrixeQTL [32]. The model was adjusted for age, gender, estimated cell proportions and three genotype-based principal components (PCs) to adjust for population sub-structure, and treated CpG-specific methylation as the outcome and SNP allele count as an explanatory variable. First, we tested each CpG site against all SNPs for both cis- and trans-SNP-CpG associations in the discovery cohort of 238 samples at diagnosis (164 CD cases and 74 controls), with adjustment for disease status in addition to other covariates. A stringent Bonferroni threshold of *P* <8.21 × 10^−14^ was used to identify mQTLs in the CD (discovery) cohort. To replicate our findings, we further tested all significant associations in the GTP cohort using the same model as in the discovery cohort, adjusting for age, gender, PTSD status, and estimated cell proportions from DNA methylation data. To identify whether these associations are specifically observed in the disease- and longitudinal datasets, we tested and compared them in the 74 healthy controls, 164 CD cases at diagnosis and at 3-year follow-up, separately. We also tested the significant blood mQTLs in a cohort of ileal samples (n=40, which is a subset of individuals from the discovery cohort) to assess tissue-specific associations. For these follow-up analyses in ileum, we restricted analysis to SNPs that had at least 2 individuals in each genotype category (AA/AB/BB) to ensure sufficient genetic variation. A statistical significance threshold of *P* < 0.05 was used for replication analyses of the disease-, longitudinal-, and tissue-associated mQTLs. We used the clump command in PLINK [15] to identify the number of independent associations for each genomic variant with more than 1 significant SNP-CpG association by using default parameters.

### Annotation of SNPs and CpG sites

We used the annovar package [33] to annotate SNPs, and the MethylationEPIC annotation file to annotate CpG sites.

### KEGG pathway enrichment analysis

We used missMethyl [34], a R/Bioconductor package, to identify KEGG database pathways that are more likely to occur in the associated CpGs than would be expected by chance. Genes with more CpG probes on the MethylationEPIC array are more likely to have differentially methylated CpGs, which could introduce potential bias when performing pathway enrichment analysis. The *gometh* function implemented in missMethyl considers the varying number of CpGs per gene by computing prior probability for each gene based on the gene length and the number of CpGs probed per gene on the array.

## List of abbreviations

(mQTL): Methylation quantitative trait loci
(IBD): Inflammatory Bowel Disease
(CD): Crohn’s Disease
(PTSD): Post-traumatic stress disorder
(GTP): Grady Trauma Project
(DNAm): DNA methylation
(GWAS): Genome-wide association studies
(SNP): Single nucleotide polymorphisms
(EWAS): Epigenome-wide association study
(MR): Mendelian Randomization
(CRP): C-reactive protein
(MAF): Minor allele frequency
(HWE): Hardy–Weinberg equilibrium
(LD): Linkage disequilibrium
(FDR): False Discovery Rate

## Declarations

### Ethics approval and consent to participate

Protocols including signed consent of all participants and/or consent of parents or legal guardians in the case of minors were approved by the IRBs of Emory University for CD RISK study supported by a research initiative grant from the Crohn’s and Colitis Foundation. The GTP study was approved by the Institutional Review Board of Emory University School of Medicine and the Grady Health Systems Research Oversight Committee. All participants provided written informed consent.

### Consent for publication

Not applicable

### Availability of data and materials

The blood DNA methylation data for all the 238 subjects (402 samples) included in this study have been deposited in the Gene Expression Omnibus (GEO) and are accessible through GEO series accession GSE112611. DNA methylation data for the 40 ileal biopsies are also deposited in GEO and can be retrieved through the accession number GSE135905.

### Authors Contributions

S.V., K.N.C., D.J.C., A.K.S. and S.K. conceived of and designed the study. J.P., D.T.O., and S. K., processed samples for methylation profiling and S.V., H.K.S., and V.K. performed the analysis with input from D.J.C. and K.N.C., K.N.C., A.K.S. and S.V., S.K., H.K.S., L.A.D., J.S.H., D.J.C., K.N.C., G.G., A.K.S. and S.K. interpreted the results and wrote the manuscript. All authors reviewed and approved the manuscript prior to submission.

## Acknowledgements

We are grateful to Anne Dodd, Jason Matthews, and all other RISK investigators for their support and helpful comments.

