## Additional file 1 for "Methylation Quantitative Trait Loci are Largely Consistent across Disease States in Crohn’s disease"

Additional file 1. Figure S1: Distribution of effect sizes (% DNA methylation change per allele) for significant SNP-CpG pairs in discovery cohort. .

Additional file 1. Figure S2: Testing the effects of previously-identified IBD SNPs and CD differentially methylated positions (DMPs) in blood and ileum.

Additional file 1. Figure S3: Boxplots for the top SNP associated to 12 CD-associated CpGs are compared between blood and ileum.

Additional file 1. Table S1: Characteristics of discovery and replication cohorts.

Additional file 2. Table S2a: The blood cis- and trans- mQTLs detected in discovery cohort with 238 pediatric samples

Additional file 1. Table S3: Genic distribution of genomic variants associated in blood cis- associations from Annovar.

Additional file 1. Table S4: Genic distribution of CpG sites in the Illumina array and cis-mQTLs

Additional file 1. Table S5: Distribution CpG island features in the Illumina array and cis-mQTLs

Additional file 3. Table S6: The blood cis-mQTLs detected in replication cohort with 780 adult African American samples

Additional file 4. Table S7: The blood cis-mQTLs tested in 164 pediatric Crohn's disease patients at baseline and follow-up

Additional file 5. Table S8: The blood mQTLs are tested in 40 ileal biopsies from pediatric cohort

Additional file 1. Table S9: The pathways enriched for the CpG sites that are negatively correlated mQTLs between blood and Ileum

Additional file 6. Table S10: The details of 69 blood mQTL associations in 238 pediatric CD patients for the known IBD genomic variants from de Lange et al., Nat Genet (2017)

Additional file 1. Table S11: The previously-identified IBD SNPs associated CPGs enriched pathways (Bonferroni < 8.21E-14)

Additional file 1. Table S12: The pathways enriched for the CpG sites that are associated with previously-identified IBD SNPs at FDR < 0.05.

Additional file 7. Table S13: The cis-mQTLs detected for 241 previously-identified IBD SNPs in discovery cohort with 238 pediatric samples, in CD cases only (n=164), in replication cohort with 780 adult African American samples, and in controls only (n=74) in blood.

Additional file 8. Table S14: The blood cis-mQTLs detected for 241 previously-identified IBD SNPs in discovery cohort are tested in all ileal biopsies (n=40), in CD cases only (n=17), and in controls only (n=23).

Additional file 9. Table S15: The cis-mQTLs detected for 1189 CD-associated CpGs in discovery cohort with 238 pediatric samples, in CD cases only (n=164), in replication cohort with 780 adult African American samples, and in controls only (n=74) in blood.

Additional file 10. Table S16: The cis-mQTLs detected for 241 previously-identified IBD SNPs in discovery cohort are tested ileal biopsies (n=40), in CD cases only (n=17), and in controls only (n=23) in ileum.

Additional file 11. Table S17: The disease-specific mQTLs showed negative effects between Cases (n=23) and Controls (n=23) in ileum

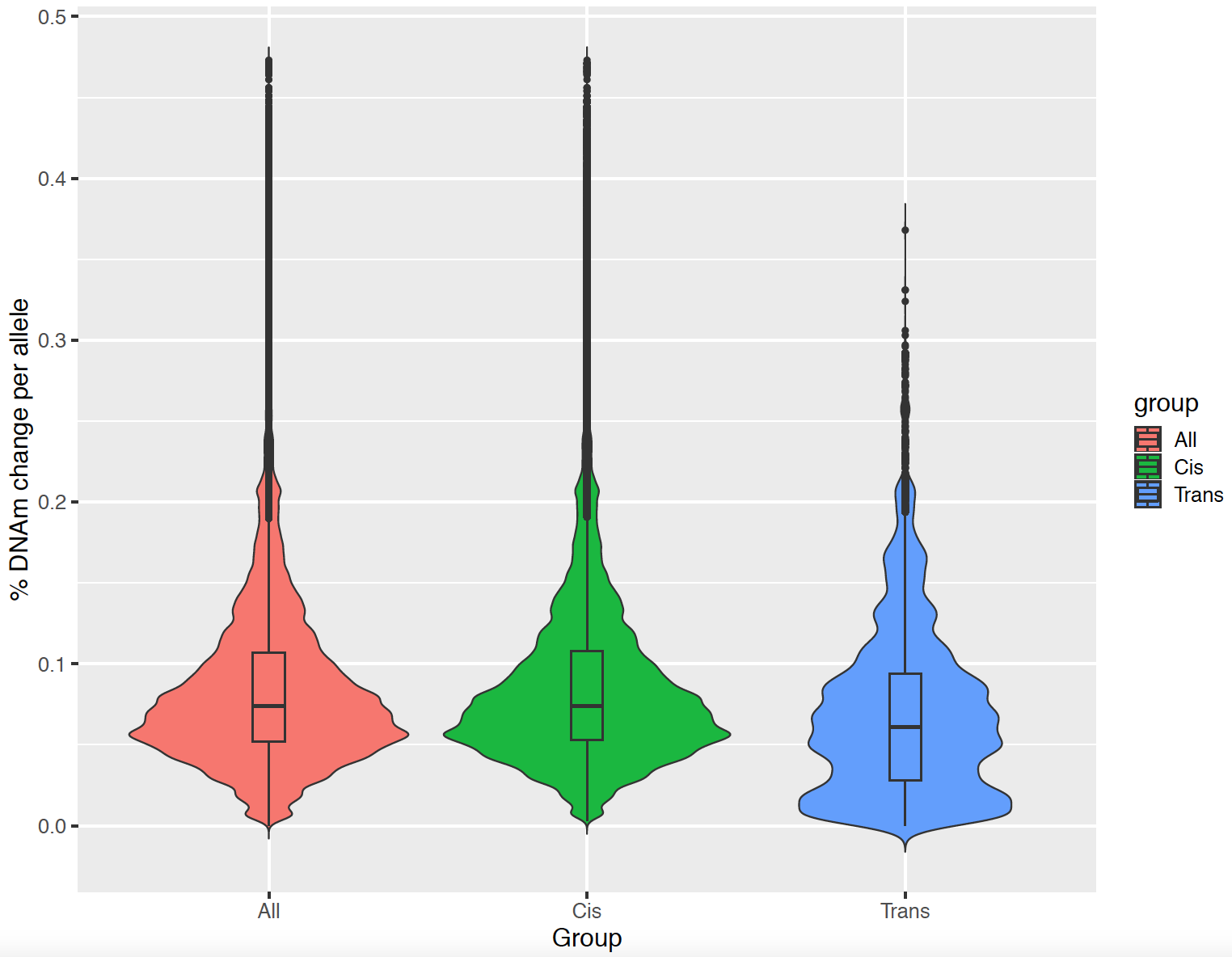

**Additional file. Figure S1:** **Distribution of effect sizes (% DNA methylation change per allele) for significant SNP-CpG pairs in discovery cohort.** The y-axis shows the DNA methylation differences associated with each minor allele of genetic variants in all SNP-CpG pairs, cis- and trans-. The average effect size for cis-mQTLs was 8.7% (IQR = 5.3 – 10.8) is significantly higher than that observed for trans-mQTLs (mean = 6.9%; IQR = 2.8 – 9.4) (Mann-Whitney-Wilcoxon Test, p < 2.2e-16).

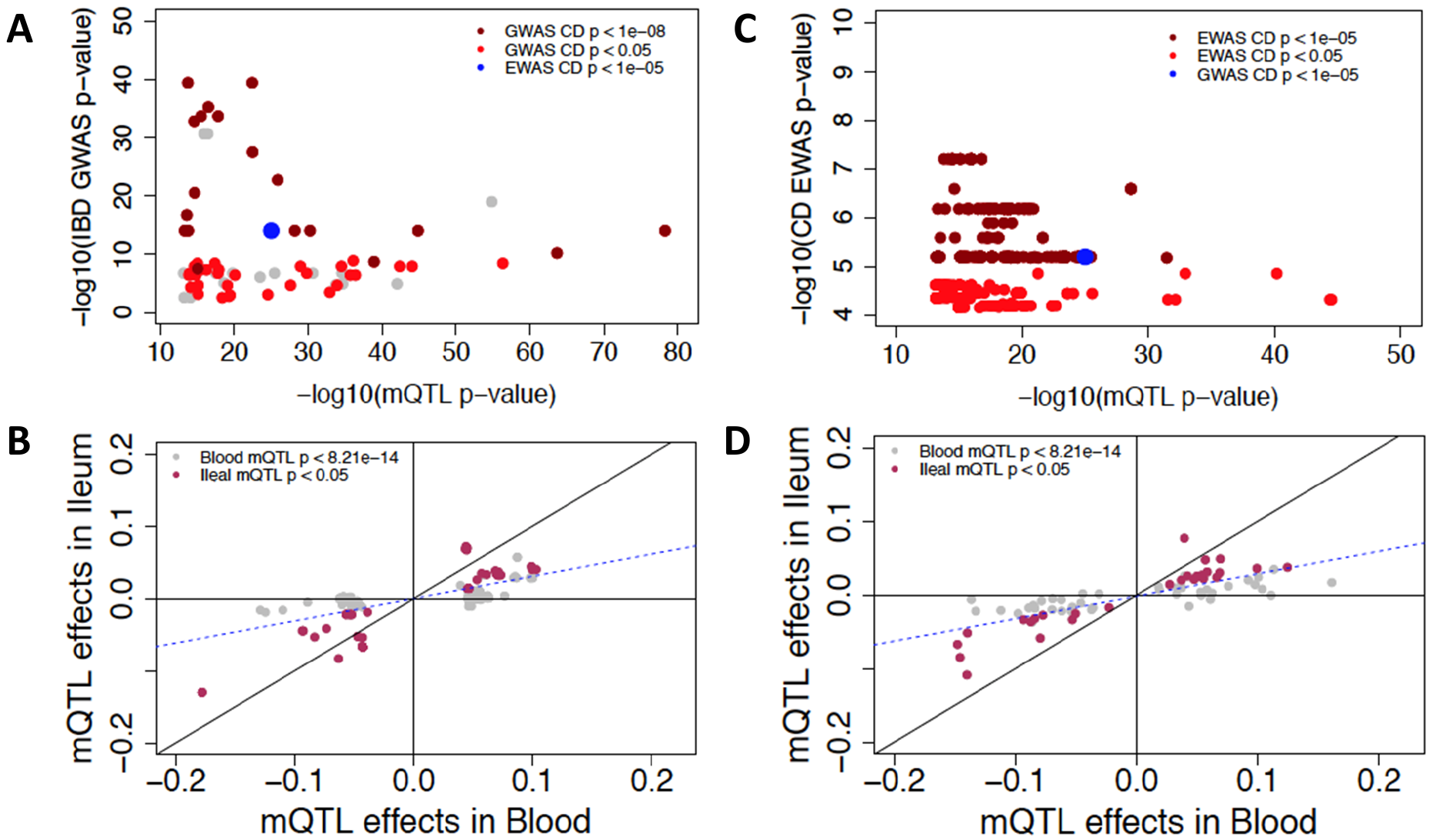

**Additional file. Figure S2: Comparison of p-values and effect sizes of previously-identified IBD SNPs and CD-associated CpG sites in blood and ileum.** (A) The y-axis shows p-values for SNPs significant in an IBD GWAS (de Lange *et al*., 2017), while the x axis shows the top mQTL associations for these SNPs in the discovery cohort. Since the discovery cohort is highly enriched with CD samples, the points are color coded based on p-values from a similar GWAS of CD (de Lange *et al*., 2017). (B) The y-axis shows EWAS p-values of CD significant CpGs from Somineni et al., (2019), while the x-axis shows the top mQTL associations for these CpGs in the discovery cohort. (C) Effect sizes of 69 SNP-CpG associations from 37 previously-identified IBD SNPs are plotted between blood and ileum. (D) Effect sizes of 565 SNP-CpG associations from 22 previously-identified IBD CpG sites are compared between blood and ileum. In both figure c and d, the maroon color indicates that the associations are Bonferroni-significant in blood (*P*=8.21x10^-14^) and nominally significant in ileum (*P* < 0.05), whereas the gray indicates the associations that are significant only in blood.

| 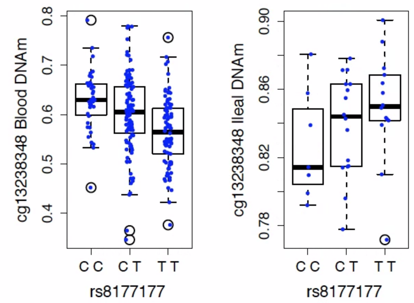 | 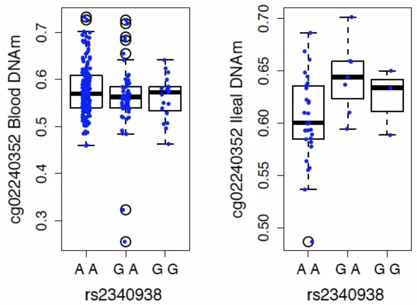 | 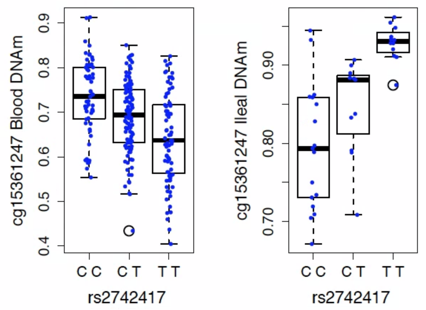 |
| --- | --- | --- |
| 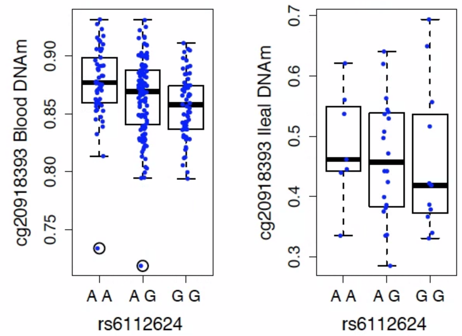 | 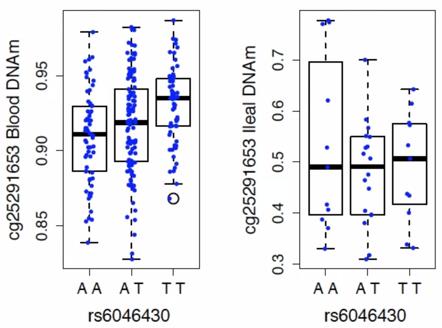 | 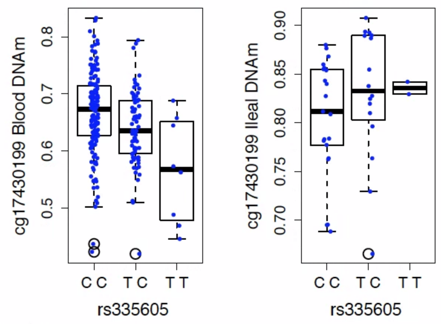 |
| 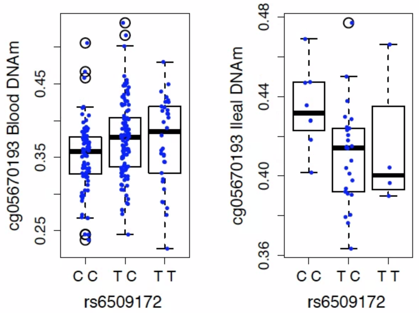 | 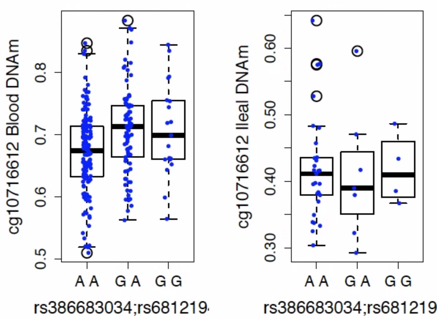 | 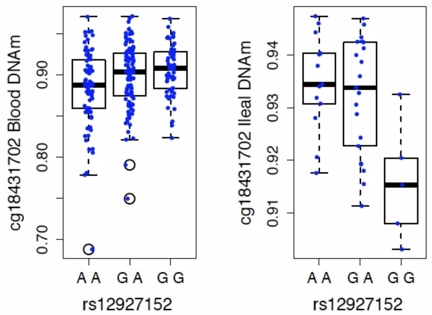 |
| 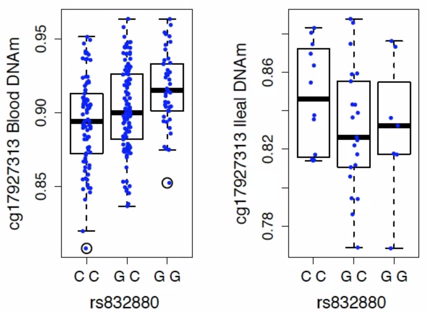 | 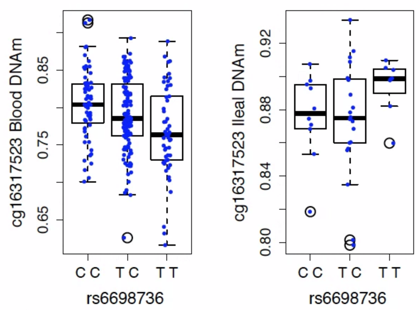 | 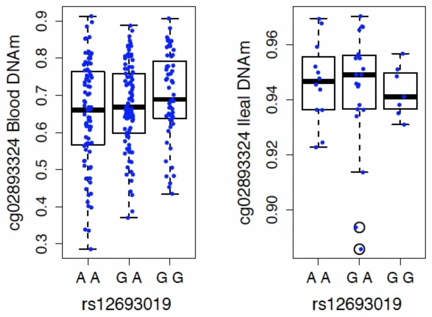 |

**Additional file. Figure S3:** Boxplots show the distribution of methylation by genotype for the top SNP associated with each of 12 CD-associated CpGs. In each panel the left plot shows this distribution in blood, and the right plot shows the distribution in ileum.

**Additional file 1. Table S1:** **Cohort description**

|  | Discovery cohort  (N=238) | Replication cohort  (N=780) | Total  (N=1018) |
| --- | --- | --- | --- |
| **Sex** | | | |
| Female | 102 (42.9%) | 556 (71.3%) | 658 (64.6%) |
| Male | 136 (57.1%) | 224 (28.7%) | 360 (35.4%) |
| **Age** | | | |
| N-Miss | 0 | 9 | 9 |
| Mean (SD) | 12.334 (3.018) | 42.263 (12.335) | 35.204 (16.733) |
| Range | 4.500 - 16.830 | 18.000 - 76.000 | 4.500 - 76.000 |
| **Race** | | | |
| African American | 40 (16.8%) | 730 (93.6%) | 770 (75.6%) |
| Caucasian | 185 (77.7%) | 25 (3.2%) | 210 (20.6%) |
| Other | 13 (5.5%) | 25 (3.2%) | 38 (3.7%) |
| **Disease status** | | | |
| Cases | 164 (68.9%) | 262 (33.6%) | 426 (41.8%) |
| Control | 74 (31.1%) | 518 (66.4%) | 592 (58.2%) |

**Additional file 1. Table S3:** **Genomic distribution of cis mQTLs identified in blood.**

|  | Total SNPs (N=3,109,862) | Cis mQTL SNPs (N=287,881) | P-value | OR (95% CI) |
| --- | --- | --- | --- | --- |
| Downstream | 20097 (0.65) | 2982 (1.04) | < 2.20E-16 | 1.6 (1.54 - 1.67) |
| Exonic | 33433 (1.08) | 5361 (1.86) | < 2.20E-16 | 1.73 (1.68 - 1.78) |
| Intergenic | 1640000 (52.74) | 127749 (44.38) | < 2.20E-16 | 0.84 (0.84 - 0.85) |
| Intronic | 1149992 (36.98) | 123635 (42.95) | < 2.20E-16 | 1.16 (1.15 - 1.17) |
| ncRNA_exonic | 11329 (0.36) | 1584 (0.55) | < 2.20E-16 | 1.51 (1.43 - 1.59) |
| ncRNA_intronic | 200858 (6.46) | 18535 (6.44) | 6.95E-01 | 1 (0.98 - 1.01) |
| ncRNA_splicing | 76 (0) | 10 (0) | 3.28E-01 | 1.42 (0.66 - 2.76) |
| Splicing | 735 (0.02) | 131 (0.05) | 1.42E-10 | 1.93 (1.59 - 2.32) |
| Upstream | 18528 (0.6) | 3056 (1.06) | < 2.20E-16 | 1.78 (1.71 - 1.85) |
| upstream;downstream | 667 (0.02) | 167 (0.06) | < 2.20E-16 | 2.7 (2.27 - 3.21) |
| UTR3 | 28373 (0.91) | 3805 (1.32) | < 2.20E-16 | 1.45 (1.4 - 1.5) |
| UTR5 | 5748 (0.18) | 850 (0.3) | < 2.20E-16 | 1.6 (1.48 - 1.72) |
| UTR5;UTR3 | 26 (0) | 16 (0.01) | 1.37E-07 | 6.65 (3.33 - 12.87) |

Table legends: downstream (1 kb region downstream of transcription end site (TES); exonic (within the exonic region); intergenic (within the intergenic region); intronic (within the intronic region); ncRNA_exonic (transcripts without coding annotation in the gene definition, within the exonic region); ncRNA_intronic (transcripts without coding annotation in the gene definition, within the intronic region); ncRNA_splicing (transcripts without coding annotation in the gene definition, within the splicing region); splicing (within 2 bp of a splicing junction); upstream (1 kb region upstream of transcription start site (TSS); UTR3 (3′ untranslated region); and UTR5 (5′ untranslated region); P-value: Fisher’s exact test p-value; OR- Odds ratio; Values in the parenthesis in each row refer % of genomic features in each category.

**Additional file 1. Table S4: Genomic distribution of all CpG sites on the Illumina array and CpG sites associated with cis-mQTLs.**

|  | Total CpGs (N=867,531) | Cis mQTL-associated CpGs (N=12,843) | P-value | OR (95% CI) |
| --- | --- | --- | --- | --- |
| Not Annotated | 250244 (28.85) | 4248 (33.08) | < 2.2E-16 | 1.22 (1.17 - 1.27) |
| 1stExon | 26433 (3.05) | 236 (1.84) | < 2.20E-16 | 0.6 (0.52 - 0.68) |
| 3'UTR | 21594 (2.49) | 227 (1.77) | 5.31E-08 | 0.70 (0.62 - 0.80) |
| 5'UTR | 73070 (8.42) | 980 (7.63) | 0.001 | 0.90 (0.84 - 0.96) |
| Body | 318165 (36.67) | 4524 (35.23) | 7.11E-04 | 0.94 (0.91 - 0.97) |
| ExonBnd | 5680 (0.65) | 47 (0.37) | 1.52E-05 | 0.56 (0.41 - 0.74) |
| TSS1500 | 107193 (12.36) | 1892 (14.73) | 2.55E-15 | 1.23 (1.17 - 1.29) |
| TSS200 | 65152 (7.51) | 689 (5.36) | < 2.2E-16 | 0.70 (0.65 - 0.75) |

Table legends: Not annotated – CpG sites were not annotated in Illumina Methylation EPIC array annotation file; 1stExon - gene’s first exon; 3′ UTR (3′ untranslated region); 5′ UTR (5′ untranslated region); Body (gene body); ExonBnd (exon boundaries), TSS1500 (200 to 1500 nucleotides (nt), upstream of transcription start site, TSS); TSS200 (up to 200 nt upstream of TSS); Values in the parenthesis in each row refer % of genomic features in each category.

**Additional file 1. Table S5: Distribution of CpG Island features for all CpG sites on the Illumina array and CpG sites associated with cis-mQTLs.**

|  | Total CpGs (N=867,531) | Cis mQTL-associated CpGs (N=12,843) | P-value | OR (95% CI) |
| --- | --- | --- | --- | --- |
| Not annotated | 489387 (56.41) | 7042 (54.83) | 3.49E-04 | 0.94(0.91 – 0.97) |
| Island | 161598 (18.63) | 1634 (12.72) | < 2.2E-16 | 0.64(0.60 – 0.67) |
| N_Shelf | 32052 (3.69) | 419 (3.26) | 9.47E-03 | 0.88(0.8 - 0.97) |
| N_Shore | 83474 (9.62) | 1757 (13.68) | < 2.20E-16 | 1.49(1.41 - 1.57) |
| S_Shelf | 29759 (3.43) | 375 (2.92) | 1.36E-03 | 0.85(0.76 - 0.94) |
| S_Shore | 71261 (8.21) | 1616 (12.58) | < 2.2E-16 | 1.61(1.52 - 1.70) |

Table legends: Not annotated – CpG sites were not annotated in Illumina Methylation EPIC array annotation file; Island – CpG Island; N_shore, S_shore - 2 kb sequences, directly up- and downstream of CpG islands are called the northern and southern shore; N_shelf and S_shelf - 2 kb sequences directly adjacent to the shores are called the northern and southern shelves, respectively; Values in the parenthesis in each row refer % of CpG island features in each category.

**Additional file 1. Table S9. Pathways enriched for genes annotated to CpG sites that show opposite-direction associations with the same SNPs between blood vs. Ileum**

| **KEGG ID** | **Pathway** | **N** | **DE** | **P.DE** | **FDR** |
| --- | --- | --- | --- | --- | --- |
| path:hsa01100 | Metabolic pathways | 1405 | 129 | 3.34E-06 | 1.11E-03 |
| path:hsa00601 | Glycosphingolipid biosynthesis - lacto and neolacto series | 27 | 7 | 1.03E-03 | 1.72E-01 |
| path:hsa04070 | Phosphatidylinositol signaling system | 99 | 17 | 3.76E-03 | 3.99E-01 |
| path:hsa04918 | Thyroid hormone synthesis | 74 | 13 | 4.79E-03 | 3.99E-01 |
| path:hsa00562 | Inositol phosphate metabolism | 74 | 12 | 1.86E-02 | 8.58E-01 |
| path:hsa04270 | Vascular smooth muscle contraction | 132 | 17 | 1.94E-02 | 8.58E-01 |
| path:hsa00514 | Other types of O-glycan biosynthesis | 22 | 5 | 2.26E-02 | 8.58E-01 |
| path:hsa04913 | Ovarian steroidogenesis | 49 | 8 | 2.64E-02 | 8.58E-01 |
| path:hsa04979 | Cholesterol metabolism | 50 | 7 | 2.99E-02 | 8.58E-01 |
| path:hsa00062 | Fatty acid elongation | 27 | 5 | 3.15E-02 | 8.58E-01 |
| path:hsa00340 | Histidine metabolism | 23 | 4 | 3.36E-02 | 8.58E-01 |
| path:hsa04912 | GnRH signaling pathway | 93 | 13 | 3.82E-02 | 8.58E-01 |
| path:hsa04064 | NF-kappa B signaling pathway | 99 | 12 | 4.40E-02 | 8.58E-01 |
| path:hsa00350 | Tyrosine metabolism | 36 | 5 | 4.96E-02 | 8.58E-01 |

Table legends: N - number of genes in the GO or KEGG term; DE - number of genes that are differentially methylated; P.DE - p-value for over-representation of the GO or KEGG term; FDR - False discovery rate

**Additional file 1. Table S11: Pathways enriched for genes annotated to CpG sites that associate with previously-identified IBD SNPs (*P* < 8.21E-14)**

|  | Pathway | N | DE | P.DE | FDR |
| --- | --- | --- | --- | --- | --- |
| path:hsa05163 | Human cytomegalovirus infection | 225 | 6 | 5.71E-06 | 1.92E-03 |
| path:hsa04931 | Insulin resistance | 108 | 4 | 7.94E-05 | 1.34E-02 |
| path:hsa04270 | Vascular smooth muscle contraction | 132 | 4 | 1.38E-04 | 1.55E-02 |
| path:hsa05161 | Hepatitis B | 162 | 4 | 2.45E-04 | 2.06E-02 |
| path:hsa04062 | Chemokine signaling pathway | 189 | 4 | 3.88E-04 | 2.62E-02 |
| path:hsa05200 | Pathways in cancer | 529 | 6 | 7.13E-04 | 3.15E-02 |
| path:hsa05212 | Pancreatic cancer | 76 | 3 | 7.35E-04 | 3.15E-02 |
| path:hsa04914 | Progesterone-mediated oocyte maturation | 95 | 3 | 9.79E-04 | 3.15E-02 |
| path:hsa04650 | Natural killer cell mediated cytotoxicity | 124 | 3 | 1.09E-03 | 3.15E-02 |
| path:hsa04912 | GnRH signaling pathway | 93 | 3 | 1.22E-03 | 3.15E-02 |
| path:hsa04933 | AGE-RAGE signaling pathway in diabetic complications | 100 | 3 | 1.25E-03 | 3.15E-02 |
| path:hsa01521 | EGFR tyrosine kinase inhibitor resistance | 78 | 3 | 1.29E-03 | 3.15E-02 |
| path:hsa04670 | Leukocyte transendothelial migration | 111 | 3 | 1.30E-03 | 3.15E-02 |
| path:hsa04666 | Fc gamma R-mediated phagocytosis | 92 | 3 | 1.33E-03 | 3.15E-02 |
| path:hsa04114 | Oocyte meiosis | 124 | 3 | 1.53E-03 | 3.15E-02 |
| path:hsa04070 | Phosphatidylinositol signaling system | 99 | 3 | 1.55E-03 | 3.15E-02 |
| path:hsa04066 | HIF-1 signaling pathway | 109 | 3 | 1.59E-03 | 3.15E-02 |
| path:hsa04928 | Parathyroid hormone synthesis, secretion and action | 106 | 3 | 2.34E-03 | 4.38E-02 |

Table legends: N - number of genes in the GO or KEGG term; DE - number of genes that are differentially methylated; P.DE - p-value for over-representation of the GO or KEGG term; FDR - False discovery rate

**Additional file 1. Table S12: Pathways enriched for genes annotated to CpGs associated with previously-identified IBD SNPs (FDR < 0.05)**

|  | Pathway | N | DE | P.DE | FDR |
| --- | --- | --- | --- | --- | --- |
| path:hsa05140 | Leishmaniasis | 74 | 6 | 3.44E-05 | 1.16E-02 |
| path:hsa04650 | Natural killer cell mediated cytotoxicity | 124 | 6 | 3.70E-04 | 4.58E-02 |
| path:hsa05200 | Pathways in cancer | 529 | 14 | 4.73E-04 | 4.58E-02 |
| path:hsa04114 | Oocyte meiosis | 124 | 6 | 7.99E-04 | 4.58E-02 |
| path:hsa04010 | MAPK signaling pathway | 295 | 10 | 8.23E-04 | 4.58E-02 |
| path:hsa04380 | Osteoclast differentiation | 127 | 6 | 8.57E-04 | 4.58E-02 |
| path:hsa04072 | Phospholipase D signaling pathway | 147 | 7 | 9.52E-04 | 4.58E-02 |
| path:hsa05235 | PD-L1 expression and PD-1 checkpoint pathway in cancer | 89 | 5 | 1.60E-03 | 6.72E-02 |
| path:hsa05321 | Inflammatory bowel disease (IBD) | 63 | 4 | 1.82E-03 | 6.81E-02 |
| path:hsa04912 | GnRH signaling pathway | 93 | 5 | 2.18E-03 | 7.12E-02 |
| path:hsa04666 | Fc gamma R-mediated phagocytosis | 92 | 5 | 2.32E-03 | 7.12E-02 |
| path:hsa05161 | Hepatitis B | 162 | 6 | 3.17E-03 | 8.07E-02 |
| path:hsa04070 | Phosphatidylinositol signaling system | 99 | 5 | 3.37E-03 | 8.07E-02 |
| path:hsa04670 | Leukocyte transendothelial migration | 111 | 5 | 3.44E-03 | 8.07E-02 |
| path:hsa05231 | Choline metabolism in cancer | 98 | 5 | 3.72E-03 | 8.07E-02 |
| path:hsa04066 | HIF-1 signaling pathway | 109 | 5 | 3.83E-03 | 8.07E-02 |
| path:hsa05163 | Human cytomegalovirus infection | 225 | 7 | 4.33E-03 | 8.59E-02 |
| path:hsa00603 | Glycosphingolipid biosynthesis - globo and isoglobo series | 15 | 2 | 6.82E-03 | 1.20E-01 |
| path:hsa04630 | JAK-STAT signaling pathway | 158 | 5 | 6.88E-03 | 1.20E-01 |
| path:hsa04060 | Cytokine-cytokine receptor interaction | 289 | 6 | 7.12E-03 | 1.20E-01 |
| path:hsa05170 | Human immunodeficiency virus 1 infection | 211 | 6 | 1.01E-02 | 1.63E-01 |
| path:hsa01521 | EGFR tyrosine kinase inhibitor resistance | 78 | 4 | 1.12E-02 | 1.71E-01 |
| path:hsa04914 | Progesterone-mediated oocyte maturation | 95 | 4 | 1.21E-02 | 1.71E-01 |
| path:hsa04020 | Calcium signaling pathway | 191 | 6 | 1.24E-02 | 1.71E-01 |
| path:hsa04727 | GABAergic synapse | 89 | 4 | 1.27E-02 | 1.71E-01 |
| path:hsa05152 | Tuberculosis | 177 | 5 | 1.39E-02 | 1.81E-01 |
| path:hsa04933 | AGE-RAGE signaling pathway in diabetic complications | 100 | 4 | 1.56E-02 | 1.95E-01 |
| path:hsa04659 | Th17 cell differentiation | 105 | 4 | 1.84E-02 | 2.09E-01 |
| path:hsa04750 | Inflammatory mediator regulation of TRP channels | 100 | 4 | 1.93E-02 | 2.09E-01 |
| path:hsa04931 | Insulin resistance | 108 | 4 | 2.05E-02 | 2.09E-01 |
| path:hsa04062 | Chemokine signaling pathway | 189 | 5 | 2.05E-02 | 2.09E-01 |
| path:hsa05131 | Shigellosis | 233 | 6 | 2.09E-02 | 2.09E-01 |
| path:hsa04015 | Rap1 signaling pathway | 210 | 6 | 2.11E-02 | 2.09E-01 |
| path:hsa04310 | Wnt signaling pathway | 160 | 5 | 2.26E-02 | 2.09E-01 |
| path:hsa00601 | Glycosphingolipid biosynthesis - lacto and neolacto series | 27 | 2 | 2.27E-02 | 2.09E-01 |
| path:hsa04370 | VEGF signaling pathway | 59 | 3 | 2.27E-02 | 2.09E-01 |
| path:hsa05145 | Toxoplasmosis | 110 | 4 | 2.30E-02 | 2.09E-01 |
| path:hsa00630 | Glyoxylate and dicarboxylate metabolism | 30 | 2 | 2.79E-02 | 2.47E-01 |
| path:hsa05031 | Amphetamine addiction | 69 | 3 | 3.07E-02 | 2.51E-01 |
| path:hsa04071 | Sphingolipid signaling pathway | 118 | 4 | 3.07E-02 | 2.51E-01 |
| path:hsa04720 | Long-term potentiation | 67 | 3 | 3.13E-02 | 2.51E-01 |
| path:hsa05135 | Yersinia infection | 120 | 4 | 3.15E-02 | 2.51E-01 |
| path:hsa04270 | Vascular smooth muscle contraction | 132 | 4 | 3.20E-02 | 2.51E-01 |
| path:hsa04935 | Growth hormone synthesis, secretion and action | 119 | 4 | 3.49E-02 | 2.66E-01 |
| path:hsa04620 | Toll-like receptor signaling pathway | 104 | 3 | 3.55E-02 | 2.66E-01 |
| path:hsa04714 | Thermogenesis | 218 | 5 | 3.81E-02 | 2.79E-01 |
| path:hsa05214 | Glioma | 75 | 3 | 4.09E-02 | 2.92E-01 |
| path:hsa04371 | Apelin signaling pathway | 137 | 4 | 4.20E-02 | 2.92E-01 |
| path:hsa04971 | Gastric acid secretion | 75 | 3 | 4.26E-02 | 2.92E-01 |
| path:hsa05212 | Pancreatic cancer | 76 | 3 | 4.33E-02 | 2.92E-01 |

Table legends: N - number of genes in the GO or KEGG term; DE - number of genes that are differentially methylated; P.DE - p-value for over-representation of the GO or KEGG term; FDR - False discovery rate
